# Population structure of UK Biobank and ancient Eurasians reveals adaptation at genes influencing blood pressure

**DOI:** 10.1101/055855

**Authors:** Kevin J. Galinsky, Po-Ru Loh, Mallick Swapan, Nick J. Patterson, Alkes L. Price

## Abstract

Analyzing genetic differences between closely related populations can be a powerful way to detect recent adaptation. The very large sample size of the UK Biobank is ideal for detecting selection using population differentiation, and enables an analysis of UK population structure at fine resolution. In analyses of 113,851 UK Biobank samples, population structure in the UK is dominated by 5 principal components (PCs) spanning 6 clusters: Northern Ireland, Scotland, northern England, southern England, and two Welsh clusters. Analyses with ancient Eurasians show that populations in the northern UK have higher levels of Steppe ancestry, and that UK population structure cannot be explained as a simple mixture of Celts and Saxons. A scan for unusual population differentiation along top PCs identified a genome-wide significant signal of selection at the coding variant rs601338 in *FUT2* (*p* = 9.16 × 10^−9^). In addition, by combining evidence of unusual differentiation within the UK with evidence from ancient Eurasians, we identified new genome-wide significant (*p* < 5 × 10^−8^) signals of recent selection at two additional loci: *CYP1A2/CSK* and *F12*. We detected strong associations to diastolic blood pressure in the UK Biobank for the variants with new selection signals at *CYP1A2/CSK* (*p* = 1.10 × 10^−19^)) and for variants with ancient Eurasian selection signals in the *ATXN2/SH2B3* locus (*p* = 8.00 × 10^−33^), implicating recent adaptation related to blood pressure.

## Introduction

Detecting signals of selection can provide biological insights into adaptations that have shaped human history^1–4^. Searching for genetic variants that are unusually differentiated between populations is a powerful way to detect recent selection^5^; this approach has been applied to detect signals of selection linked to lactase resistance^6,7^, fatty acid decomposition^8^, hypoxia response^9–11^, malaria resistance^12–14^, and other traits and diseases^15–18^.

Leveraging population differentiation to detect selection is particularly powerful when analyzing closely related subpopulations with large sample sizes^19^. Here, we analyze 113,851 samples of UK ancestry from the UK Biobank (see URLs) in conjunction with recently published People of the British Isles (PoBI)^20^ and ancient DNA^21–24^ data sets to draw inferences about population structure and recent selection. We employ a recently developed selection statistic that detects unusual population differentiation along continuous principal components (PCs) instead of between discrete subpopulations^25^, and combine our results with independent results from ancient Eurasians^23^. We detect three new signals of selection, and show that genetic variants with both new and previously reported^23^ signals of selection are strongly associated to diastolic blood pressure in UK Biobank samples.

## Results

### Population Structure in the UK Biobank

We restricted our analyses of population structure to 113,851 UK Biobank samples of UK ancestry and 202,486 SNPs after quality control (QC) filtering and linkage disequilibrium (LD) pruning (see Online Methods). We ran principal components analysis (PCA) on this data, using our FastPCA implementation^25^ (see URLs). We determined that the top 5 PCs represent geographic population structure (Figure 1), by visually examining plots of the top 10 PCs (SupplementaryFigure 1), observing that the eigenvalues for the top 5 PCs were above background levels, and that the eigenvectors were correlated with birth coordinate (SupplementaryTable 1). The eigenvalue for PC1 was 20.99, which corresponds to the eigenvalue that would be expected at this sample size for two discrete subpopulations of equal size with an *F_ST_* of 1.76 × 10^−4^ (SupplementaryTable 1).

**Figure 1.**
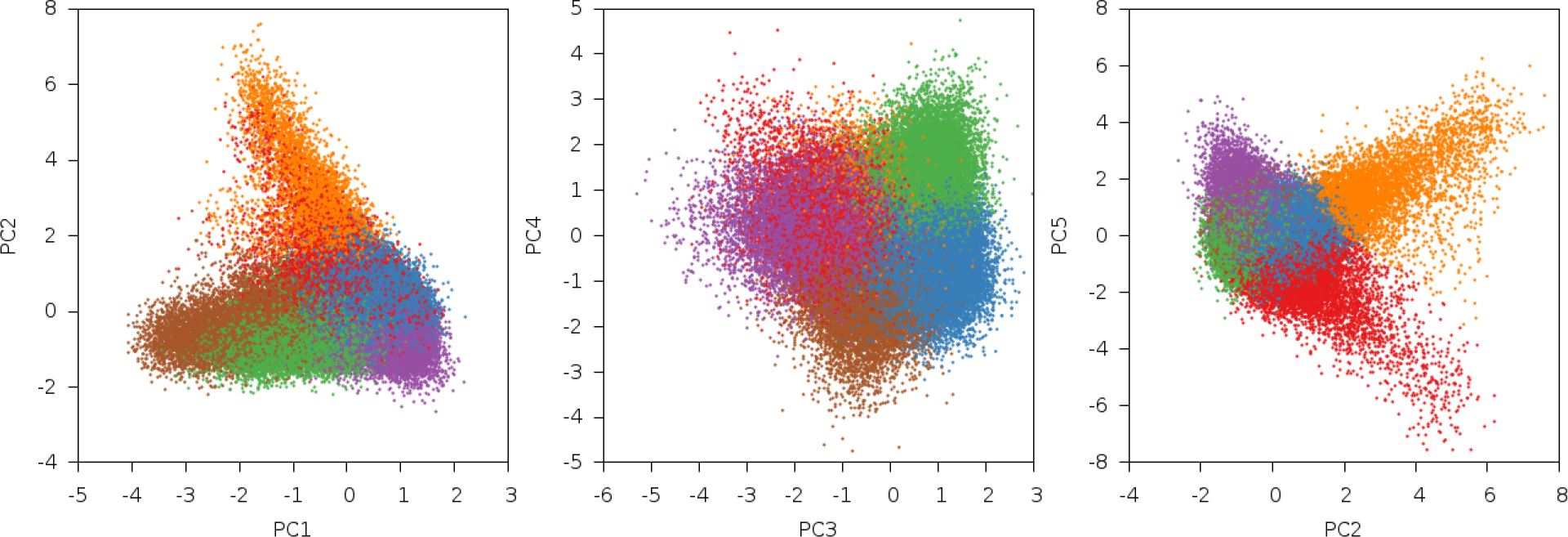
Results of PCA with *k*-means clustering. The top 5 PCs in UK Biobank data are displayed. Samples were clustered using these PCs into 6 clusters with *k*-means clustering (see Table 1). PC5 is plotted against PC2, because PC5 primarily separated the orange and red clusters, which were separated from the other clusters by PC2.

We ran *k*-means clustering on these 5 PCs to partition the samples into 6 clusters, since *K* PCs can differentiate *K* + 1 populations (Figure 1, Table 1, SupplementaryFigure 2). To identify the populations underlying the 6 clusters, we projected the PoBI dataset^20^, comprising 2,039 samples from 30 regions of the UK, onto the UK Biobank PCs (Figure 2, SupplementaryFigure 3). The individuals in the PoBI study were from rural areas of the UK and had all four grandparents born within 80 km of each other, allowing a glimpse into the genetics of the UK before the increase in mobility of the 20^th^ century. We selected representative PoBI sample regions that best aligned with the 6 UK Biobank clusters by comparing centroids of each projected population region with those from the UK Biobank clusters via visual inspection (see Online Methods, Table 1). The largest cluster represented southern England, three clusters represented different regions in the northern UK (northern England, Northern Ireland and Scotland) and two clusters represented north and south Wales. The PCs separated the six UK clusters along two general geographical axes: a north-south axis and a Welsh-specific axis. PC1 and PC3 both separated individuals on north-south axes of variation, with southern England on one end and one of the northern UK clusters on the other. PC2 separated the Welsh clusters from the rest of the UK. PC4 separated the Scotland cluster from the Northern Ireland cluster. PC5 separated the north Wales and south Wales (also known as Pembrokeshire) clusters from each other.

**Figure 2.**
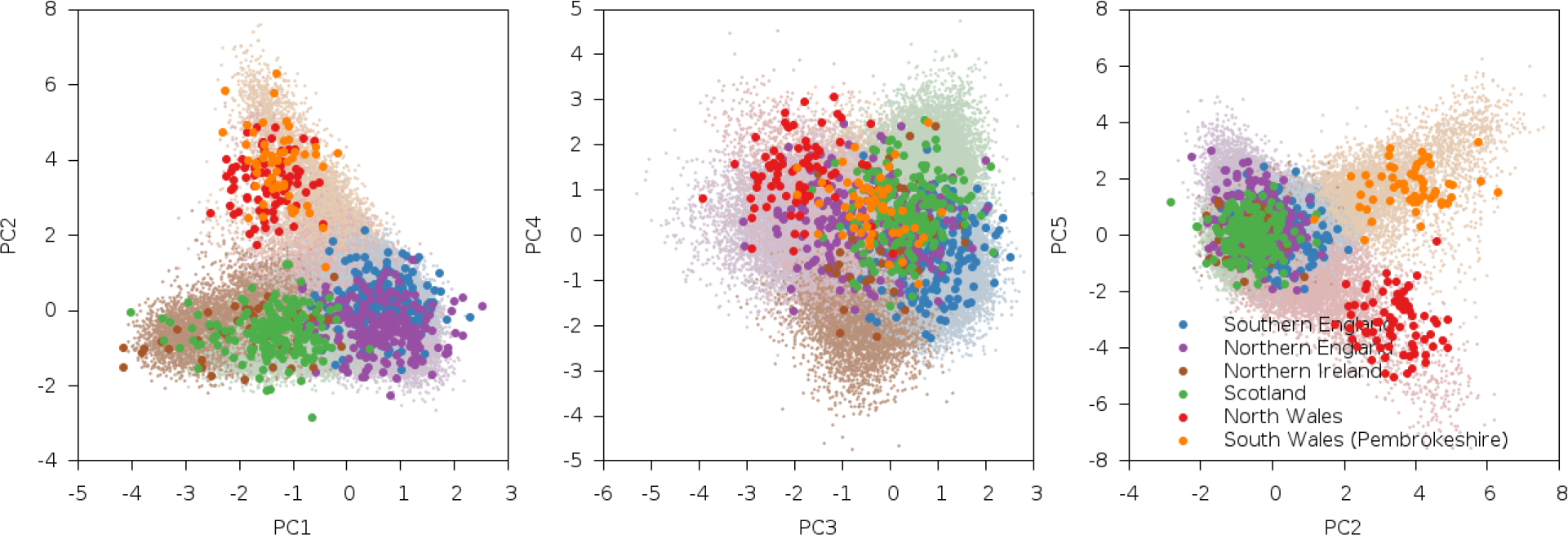
Results of PCA with projection of PoBI samples. The top 5 PCs in UK Biobank data are displayed with PoBI samples projected onto these PCs. PoBI populations which visually best matched the clusters from *k*-means clustering were used to assign names to the six clusters (Table 1).

**Table 1.** Correspondence between UK Biobank clusters and PoBI populations. We report the PoBI population that most closely corresponds to each UK Biobank cluster (see main text).

| Color | Count | Cluster Name | PoBI Populations |
| --- | --- | --- | --- |
| Blue | 41,494 | Southern England | Hampshire, Devon, Norfolk |
| Purple | 19,452 | Northern England | Yorkshire, Lancashire |
| Brown | 12,895 | Northern Ireland | Northern Ireland |
| Green | 21,215 | Scotland | Argyll and Bute, Banff and Buchan, Orkney |
| Red | 14,190 | North Wales | North Wales |
| Orange | 4,605 | South Wales / Pembrokeshire | North Pembrokeshire, South Pembrokeshire |

We next analyzed UK Biobank population structure in conjunction with ancient DNA samples. Modern European populations are known to have descended from three ancestral populations: Steppe, Mesolithic Europeans and Neolithic farmers^21,22^. We projected ancient samples from these three populations as well as ancient Saxon samples^24^ onto the UK Biobank PCs (Figure 3, SupplementaryFigure 4, see Online Methods). These populations were primarily differentiated along PC1 and PC3, indicating higher levels of Steppe ancestry in northern UK populations.

**Figure 3.**
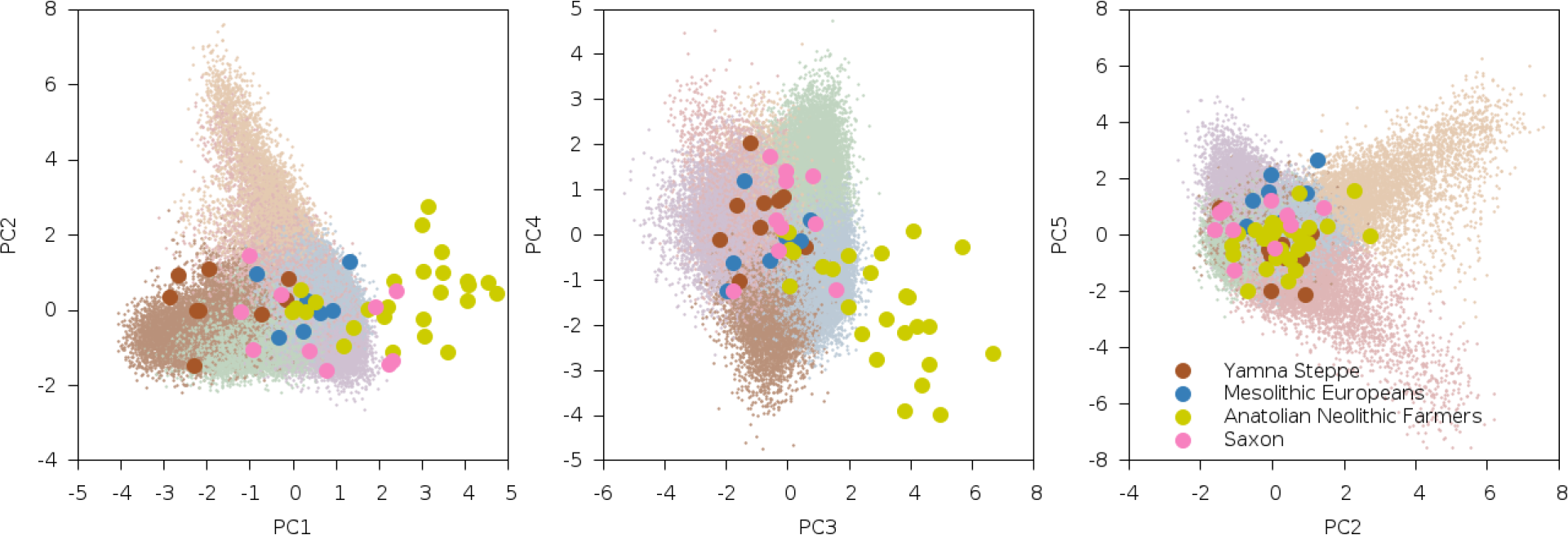
Results of PCA with projection of ancient samples. The top 5 PCs in UK Biobank data are displayed with ancient samples projected onto these PCs.

Additionally, the lack of any ancient sample correlation with PC2 suggests that Welsh populations are not differentially admixed with any ancient population in our data set, and likely underwent Welsh-specific genetic drift. We confirmed these findings by projecting pan-European POPRES^26^ samples onto the UK Biobank PCs (see Online Methods, SupplementaryFigure 5) noting that of the continental European populations, Russians (who have the most Steppe ancestry) lie on one side and Spanish and Italians (who have least)^22^ lie on the other side along PC1 and PC3, and that none of the continental European populations projected onto the same regions as the Welsh on PC2 and PC5.

In addition to the impact of ancient Eurasian populations, we know that the genetics of the UK has been strongly impacted by Anglo-Saxon migrations since the Iron Age^24^, with the Angles arriving in eastern England and the Saxons in southern England. The Anglo-Saxons interbred with the native Celts, which explains much of the genetic landscape in the UK. We analyzed a variety of samples from Celtic (Scotland and Wales) and Anglo-Saxon (southern and eastern England) populations from modern Britain in conjunction with the PoBI samples^20^ and 10 ancient Saxon samples from eastern England^24^ in order to assess the relative amounts of Steppe ancestry. We computed *f*_4_ statistics^27^ of the form *f*_4_(*Steppe*, *Neolithic Farmer*; *Pop*1, *Pop*2), where *Steppe* and *Neolithic Farmer* populations are from ref. ^21,22^, *Pop1* is either a modern Celtic or ancient Saxon population and *Pop2* is a modern Anglo-Saxon population (Table 2, SupplementaryTable 2). This statistic is sensitive to Steppe ancestry with positive values indicating more Steppe ancestry in *Pop1* than *Pop2*. We consistently obtained significantly positive *f*_4_ statistics, implying that both the modern Celtic samples and the ancient Saxon samples have more Steppe ancestry than the modern Anglo-Saxon samples from southern and eastern England. This indicates that southern and eastern England is not exclusively a genetic mix of Celts and Saxons. There are a variety of possible explanations, but one is that the present genetic structure of Britain, while subtle, is quite old, and that southern England in Roman times already had less Steppe ancestry than Wales and Scotland.

**Table 2.** Results of *f*_4_ statistics in ancient and modern British samples. We report *f*_4_ statistics of of the form *f*_4_(*Steppe*, *Neolithic Farmer*; *Pop*1, *Pop*2), representing a z-score with positive values indicating more Steppe ancestry in *Popl* than *Pop2*. Samples for *Popl* were either modern Celtic (Scotland and Wales) or ancient Saxon. Samples for *Pop2* were modern Anglo-Saxon (southern and eastern England).

|  |  | ***Pop2*** |  |  |
| --- | --- | --- | --- | --- |
| **Grouping** | ***Pop1*** | Hampshire | Devon | Norfolk |
| Ancient | Saxon | 2.543 | 3.732 | 5.118 |
| Scotland | Argyll and Bute | 3.323 | 6.223 | 9.560 |
| North Wales | North Wales | 1.918 | 5.239 | 8.490 |
| South Wales | North Pembrokeshire | 1.759 | 4.430 | 7.124 |

### Signals of Natural Selection

We searched for signals of selection using a recently developed selection statistic that detects unusual population differentiation along continuous PCs^25^. Notably, this statistic is able to detect selection signals at genome-wide significance. We analyzed the top 5 UK Biobank PCs (which were computed using LD-pruned SNPs), and computed selection statistics at 510,665 SNPs, reflecting the set of SNPs after QC but before LD-pruning (see Online Methods). The Manhattan plot for PC1 is reported in Figure 4, with additional plots in SupplementaryFigure 6. We detected genome-wide significant signals of selection at *FUT2* and at several loci with widely known signals of selection (Table 3). Loci with suggestive signals of selection (*p* < 10^−6^) are reported in SupplementaryTable 3. *FUT2* has also previously been reported as a target of natural selection^28,29^, although those results focused on frequency differences between highly diverged continental populations whereas our results implicate much more recent selection. *FUT2* encodes fucosyltransferase 2, an enzyme that affects the Lewis blood group. The SNP with the most significant *p*-value, rs601338, is a coding variant where the variant rs601338*G encodes the secretor allele and the rs601338*A variant encodes the nonsecretor allele, which protects against the Norwalk norovirus^30,31^. This SNP also affects the progression of HIV infection^32^, and is associated with vitamin B_12_ levels^33^, Crohn’s disease^34^, celiac disease and inflammatory bowel disease^35^, possibly due to changes in gut microbiome energy metabolism^36^. rs601338*A is more common in northern UK samples (SupplementaryTable 4). Similar allele frequency patterns were also observed in GERA^37^ and PoBI^20^ samples at rs492602 and rs676388 (SupplementaryTable 4), two linked SNPs in *FUT2* whose allele frequencies vary on a north-south axis in UK Biobank data. rs492602 and rs676388 were suggestively significant (p < 1.00 × 10 ^−6^) but not genome-wide-significant in tests for selection using the GERA data set (SupplementaryTable 5), emphasizing the advantage of analyzing more closely related subpopulations in very large sample sizes in the UK Biobank data set. These three SNPs were also significant when analyzing the 6 UK Biobank clusters described above using a test for selection based on unusual differentiation between discrete subpopulations (SupplementaryTable 6).

**Figure 4.**
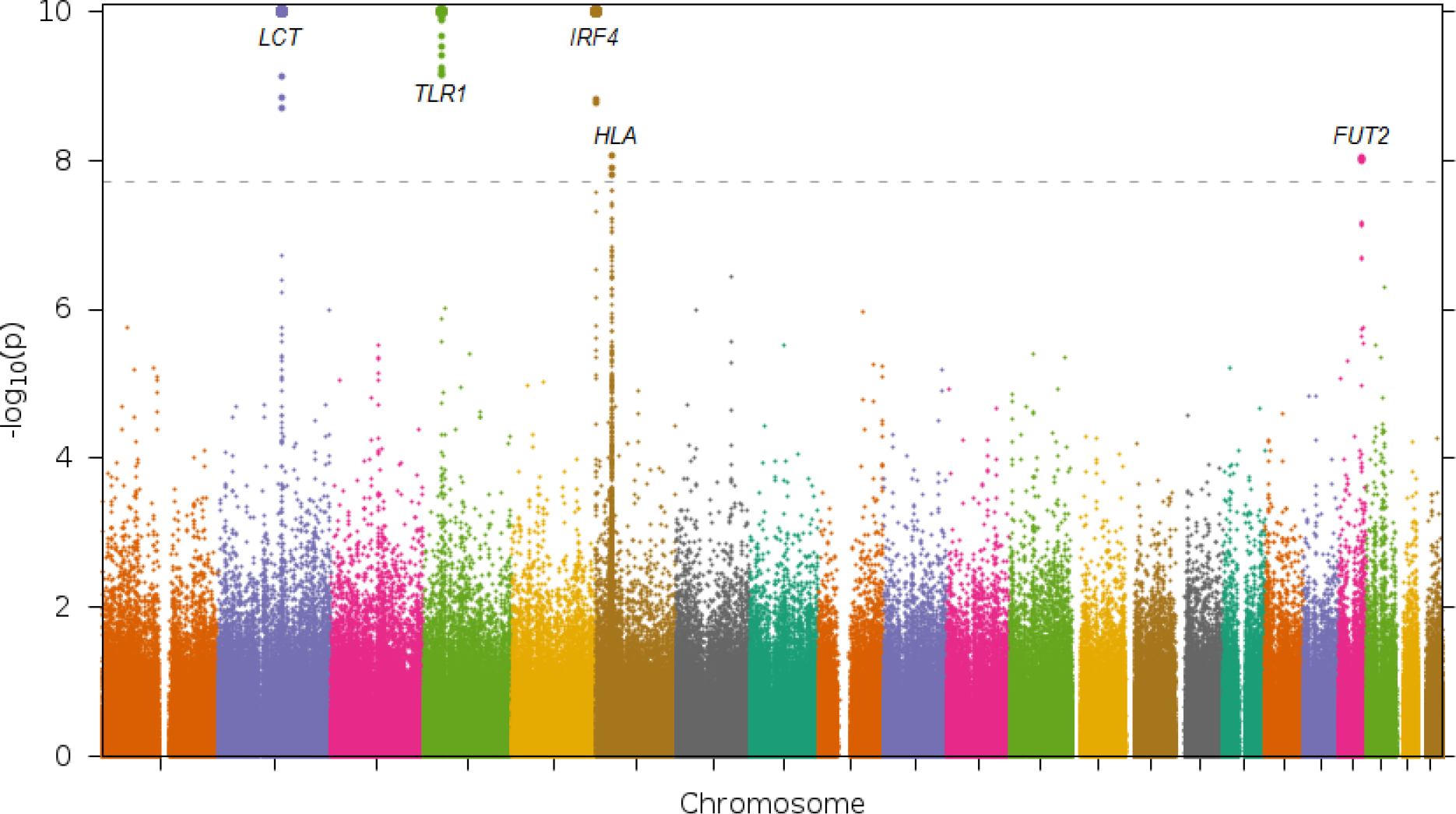
Selection statistics for UK Biobank along PC1. A Manhattan plot with – log_10_(*p*) values is displayed. Values above the significance threshold (dotted line, *p* = 1.96 × 10^−8^, *α* = 0.05 after correcting for 5 PCs and 510,665 SNPs) are displayed as larger points and are labeled with the locus they correspond to (see Table 3). – log_1o_(*p*) values larger than 10 are truncated at 10 for easier visualization and are displayed as even larger points.

**Table 3.** Top signals of selection for UK Biobank along PC1-PC5. We report the top signal of natural selection for each locus reaching genome-wide significance (*p* < 1.96 × 10^−8^) along any of the top five PCs. Neighboring SNPs <1Mb apart with genome-wide significant signals were grouped together into a single locus.

| Locus | Chromosome | Position (Mb) | PC | Top SNP | *p*-value |
| --- | --- | --- | --- | --- | --- |
| **LCT**<sup>6</sup> | 2 | 134.9 - 137.2 | 1 | rs7570971 | $3.96 \times 10^{-15}$ |
| **TLR1**<sup>68</sup> | 4 | 38.8 - 38.9 | 1 | rs4833095 | $7.96 \times 10^{-15}$ |
| | | | 2 | | $1.27 \times 10^{-8}$ |
| | | | 3 | | $7.89 \times 10^{-9}$ |
| | | | 4 | | $1.54 \times 10^{-11}$ |
| **IRF4**<sup>69,70</sup> | 6 | 0.4 - 0.5 | 1 | rs62389423 | $2.31 \times 10^{-43}$ |
| **HLA**<sup>71</sup> | 6 | 31.1 - 32.9 | 1 | rs9366778 | $8.45 \times 10^{-9}$ |
| **FUT2** | 19 | 49.2 - 49.2 | 1 | rs601338 | $9.16 \times 10^{-9}$ |

To detect additional signals of selection, we combined our PC-based selection statistics from the UK Biobank data with a previously described selection statistic that detects unusual allele frequency differences after the admixture of ancient Eurasian populations by identifying SNPs whose allele frequencies are inconsistent with admixture proportions inferred from genome-wide data^23^. For each of PC1-PC5 in UK Biobank, we summed our chi-square (1 d.o.f.) selection statistics for that PC with the chi-square (4 d.o.f.) selection statistics from ref. 23 to produce chi-square (5 d.o.f.) statistics that combine these independent signals (see Online Methods). We confirmed the independence of the two selection statistics by checking that the combined statistics were not inflated, as well as by examining the correlations between the two selection statistics (SupplementaryTable 7). We looked for signals that were genome-wide significant in the combined selection statistic but not in either of the constituent UK Biobank or ancient Eurasian selection statistics. Results are reported in Table 4.

**Table 4.** Top signals of selection for combined selection statistics. We report the top selection statistic for each locus reaching genome-wide significance, restricting to loci that were not genome-wide significant in either the UK Biobank selection statistic or the ancient Eurasian selection statistics. Neighboring SNPs <1Mb apart with genome-wide significant signals were grouped together into a single locus.

| Locus | Chr | Position (Mb) | PC | Top SNP | Combined $p$-value | UK Biobank $p$-value | Ancient Eurasian $p$-value |
| --- | --- | --- | --- | --- | --- | --- | --- |
| ***F12*** | 5 | 33.9 - 34.0 | 2 | rs2545801 | $1.79 \times 10^{-9}$ | $4.36 \times 10^{-4}$ | $5.35 \times 10^{-8}$ |
| ***CYP1A2 / CSK*** | 15 | 75.0 - 75.1 | 2 | rs1378942 | $4.65 \times 10^{-8}$ | $1.05 \times 10^{-2}$ | $1.08 \times 10^{-7}$ |

We detected new genome-wide significant signals of selection at the *F12* and *CYP1A2/CSK* loci. We are not currently aware of previous evidence of selection at *F12*. *F12* codes for coagulation factor XII, a protein involved in blood clotting^38^. The SNP at the *F12* locus, rs2545801 was suggestively significant in the ancient Eurasian analysis (*p* = 5.35 × 10^−8^), and combining it with the UK Biobank selection statistic on PC2 produced a genome-wide significant signal. This SNP has been associated with activated partial thromboplastin time, a measure of blood clotting speed where shorter time is a risk factor for strokes^39^. An additional significant SNP at *F12*, rs2731672, affects expression of *F12* in liver^40^ and is associated with plasma levels of factor XII^41^. The *CYP1A2/CSK* locus has previously been reported as a target of natural selection when comparing inter-continental allele and haplotype frequencies^42,43^, but our results implicate much more recent selection. The two detected SNPs at this locus are in strong LD (*r*^2^ = 0.858). The top SNP, rs1378942, is in an intron in the *CSK* gene. This SNP has greatly varying allele frequency across continents^43^, is associated with blood pressure^44,45^ and systemic sclerosis (an autoimmune disease affecting connective tissue)^46^. The second SNP, rs2472304 in *CYP1A2*, is associated with esophageal cancer^47^, caffeine consumption^48^ and may mediate the protective effect of caffeine on Parkinson’s disease^49^.

We tested SNPs with genome-wide significant signals of selection in the constituent UK Biobank or ancient Eurasian scans or the combined scan for association with 15 phenotypes in the UK Biobank data set, using the top 5 PCs as covariates (SupplementaryTable 8, see Online Methods). The top SNP at *F12* (rs2545801) was associated with height (*p* = 4.8 × 10^−11^), and the top SNP at *CYP1A2/CSK* (rs1378942) was associated with diastolic blood pressure (DBP) (*p* = 3.6 × 10^−19^) and hypertension (*p* = 4.8 × 10^−9^), consistent with previous findings^50^. We detected additional associations with DBP (*p* = 8.00 × 10^−33^) and hypertension (*p* = 1.30 × 10^−9^) at the *ATXN2/SH2B3* locus which was reported as under selection in the ancient Eurasian scan. The top SNP in *ATXN2/SH2B3,* rs3184504, is known to be associated with blood pressure^51^. We note that PC1 and PC3 were strongly associated with height in the UK Biobank data set, and PC3 and PC4 were associated with DBP(SupplementaryTable 9). *GRK4*^52^, *AGT*^52^ and *ATP1A1*^14^ have also been reported to be under selection and to be associated with DBP or hypertension. None of the SNPs in *GRK4* or *ATP1A1* were found to be under selection or associated with DBP or hypertension in our analyses. The *AGT* SNP rs699 was associated with DBP (*p* = 7.2 × 10^−10^) and nominally associated to hypertension (*p* = 4.8 × 10^−4^), although it did not produce a significant signal of selection in our analyses.

## Discussion

In this study, we used PCA to analyze the population structure of a large UK cohort (*N* = 113,851). We detected 5 PCs representing geographic population structure that partitioned this cohort into six subpopulation clusters. Projecting ancient samples onto these PCs revealed greater Steppe ancestry in northern UK samples. No ancient samples were found to vary along the Welsh-specific axis, suggesting that the Welsh populations differ from the rest of the UK due to drift and not different levels of admixture. We also determined that UK population structure cannot be explained as a simple mixture of Celts and Saxons.

We leveraged the subtle population structure and large sample size of the UK Biobank data set to detect signals of natural selection. We determined that the rs601338*A allele of *FUT2* was more common in northern UK samples, suggesting that pathogens may have exerted selective pressure in those populations. Combining a selection statistic that detects selection via population differentiation within the UK with a separate statistic that detects selection since ancient population admixture in Europe, we were able to detect selection at two additional loci, *F12* and *CYP1A2/CSK*. We additionally found associations to diastolic blood pressure at *CYP1A2/CSK*and at the *ATXN2/SH2B3* locus implicated in a previous selection scan.

We conclude by noting three limitations in our work. First, we employed PCA, a widely used method for analyzing population structure^25,53,54^, but haplotype-based methods such as fineSTRUCTURE may be more powerful^20,55,56^; recent advances in computationally efficient phasing^57,58^ increase the prospects for applying such methods to biobank scale data. Second, we employed methods designed to detect selection at individual loci, but did not employ methods to detect polygenic selection^59–63^; our observation that top PCs were correlated with height and DBP in the UK Biobank data set, which could potentially be consistent with the action of polygenic selection on these traits, motivates further analyses of possible polygenic selection. Finally, the PC-based test for selection that we employed assumes that allele frequencies vary linearly along a PC. The spatial ancestry analysis (SPA) method^64–66^ allows for a logistic relationship between allele frequency and ancestry, and is not constrained by this limitation. However, the advantage of the PC-based test for selection is that it allows for the detection of genome-wide significant signals, a key consideration in genome scans for selection.

## Online Methods

### UK Biobank data set

The UK Biobank phase 1 data release contains 847,131 SNPs and 152,729 samples. We removed SNPs that were multi-allelic, had a genotyping rate less than 99%, or had minor allele frequency (MAF) less than 1%. We also removed samples with non-British ancestry as well as samples with a genotyping rate less than 98%. This left 510,665 SNPs and 118,650 samples, a data set that we call “QC*.” Using PLINK2^67^ (see URLs), we removed SNPs not in Hardy-Weinberg equilibrium (*p* < 10^−6^), and we LD-pruned SNPs to have r^2^ < 0.2. We then generated a genetic relationship matrix (GRM) and removed one of each any pair of samples with relatedness greater than 0.05. This data set, which we call “LD,” contained 210,113 SNPs and 113,851 samples. Taking the full set of SNPs from the QC* data set and the set of unrelated samples from the LD data set produces the final “QC” dataset.

### PoBI and POPRES data sets

The 2,039 UK PoBI samples were a subset of the 4,371 samples collected as part of the PoBI project^20^. The 2,039 samples were a subset of the 2,886 samples genotyped on the Illumina Human 1.2M-Duo genotyping chip, with 2,510 samples passing QC procedures and 2,039 samples with all four grandparents born within 80km of each other. We also examined 2,988 European POPRES samples from the LOLIPOP and CoLaus collections^26^. These samples were genotyped on the Affymetrix GeneChip 500K Array.

### Ancient DNA data sets

Ancient DNA was gathered from several regions. 9 Steppe samples were collected from the Yamna oblast in Russia^22^, 7 west-European hunter-gatherers from Loschbour^21^, 26 Neolithic farmer samples from the Anatolian region^22^, and 10 Saxon samples from three sites in the UK^24^. DNA was extracted from bone tissue, PCR amplified and then purified using a hybrid capture approach^22–24^. The resulting DNA was sequenced on Illumina MiSeq, HiSeq or NextSeq platforms. Sequenced reads were aligned to the human genome using BWA and called SNPs were intersected with the SNPs found on the Human Origins Array^27^.

### PCA

We ran PCA on the UK Biobank LD dataset using the FastPCA software in EIGENSOFT^25^ (see URLs). We identified several artifactual PCs that were dominated by regions of long-range LD (Supplementary Figure 7). Removing loci with significant or suggestive selection signals (SupplementaryTable 10) along with their flanking 1Mb regions from the LD data set and rerunning PCA eliminated these artifactual PCs (SupplementaryFigure 1). We refer to the resulting data set with 202,486 SNPs and 113,851 samples as the “PC” dataset.

### PC Projection

We projected PoBI^20^ (642,288 SNPs, 2,039 samples from 30 populations), POPRES^26^ (453,442 SNPs, 4,079 samples from 60 populations) and ancient DNA^22,23^ (159,588 SNPs, 52 samples from 4 populations) samples onto the UK Biobank PCs via PC projection^53^. The SNPs in the UK Biobank QC data set were intersected with those in the projected data set and A/T and C/G SNPs were removed due to strand ambiguity (75,254, 37,593 and 24,467 SNPs for PoBI, POPRES and ancient DNA, respectively). The intersected set of SNPs was stringently LD-pruned for r^2^ < 0.05 using PLINK2^67^ (see URLs) (leaving 27,769, 20,914 and 15,722 SNPs respectively). SNP weights were computed for the intersected set of SNPs and these weights were then used to project the new samples onto the UK Biobank PCs^53^.

### PCA-based selection statistic

PCA is equivalent to the singular value decomposition (***X*** = ***U*Σ*V***^*T*^) where ***X*** is the normalized genomic matrix, ***U*** is the matrix of left singular vectors, ***V*** is the matrix of right singular vectors, and **Σ** is a diagonal matrix of singular values. The singular values are related to the eigenvalues of the genetic relationship matrix (GRM) by the relationship Λ = Σ^2^/*M*, where *M* is the number of SNPs used to compute the GRM ***X***^*T*^***X***/*M*. The matrix ***U*** has the properties ***U***^*T*^***U*** = ***I*** and ***U*** = *X**v*****Σ**^−1^. By the central limit theorem, the elements of ***U*** follow a normal distribution and after rescaling by *M* they follow a chi-square (1 d.o.f.) distribution. In other words, the statistic 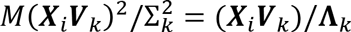 for the i^th^ SNP at the k^th^ PC follows a chi-square (1 d.o.f.) distribution^25^. One benefit of this statistic is that the PCs can be generated on one set of SNPs (here we used the PC dataset described earlier) and the selection statistic can be calculated on another set of SNPs (we used the QC dataset).

Signals of selection were clustered by considering all SNPs for which the p-value along at least one PC was less than an initial threshold (which we set at 10^−6^) and clustering together SNPs within 1Mb. We defined genome-wide significant loci based on clusters that contained at least one SNP with a *p*-value smaller than the genome-wide significance threshold. Since we analyzed 5 PCs and 510,665 SNPs, the genome-wide significance threshold was 0.05/(5 × 510,665) = 1.96 × 10^−8^. We defined suggestive loci based on clusters with at least two SNPs crossing the initial threshold (but none crossing the genome-wide significance threshold).

### Combined selection statistic

We intersected the chi-square (4 d.o.f.) ancient Eurasian selection statistics for 1,004,613 SNPs from Mathieson *et al.*^23^ with the PC-based chi-square (1 d.o.f.) UK Biobank selection statistics for 510,665 QC SNPs, producing a list of 115,066 SNPs. For each SNP and each PC, we added the ancient Eurasian selection statistics to the UK Biobank selection statistics for that PC, producing chi-square (5 d.o.f.) statistics which we corrected using genomic control.

### Association tests

Association analyses were performed using PLINK2^67^ with the top 5 PC as covariates using the “––linear” or “––logistic” flags.

## Acknowledgments

We thank Iain Mathieson and David Reich for helpful discussions and Stephan Schiffels for technical assistance with Saxon samples. This research was conducted using the UK Biobank Resource and was funded by NIH grant R01 HG006399.

## URLs

UK Biobank: http://www.ukbiobank.ac.uk/

EIGENSOFT v6.1.1 (FastPCA and PC-based selection statistic): http://www.hsph.harvard.edu/alkes-price/software/

PLINK2: https://www.cog-genomics.org/plink2

## Supplementary Figures

**Supplementary Figure 1 Results of PCA after removing long-range LD regions**

Regions with high SNP weights from the first PCA run were removed and PCA was run on the remainder of the genome (see Online Methods). The resulting PCs are no longer influenced by long-range LD regions. A visual inspecion suggests that PC1-PC5 have interesting population structure while PC6-PC10 do not.

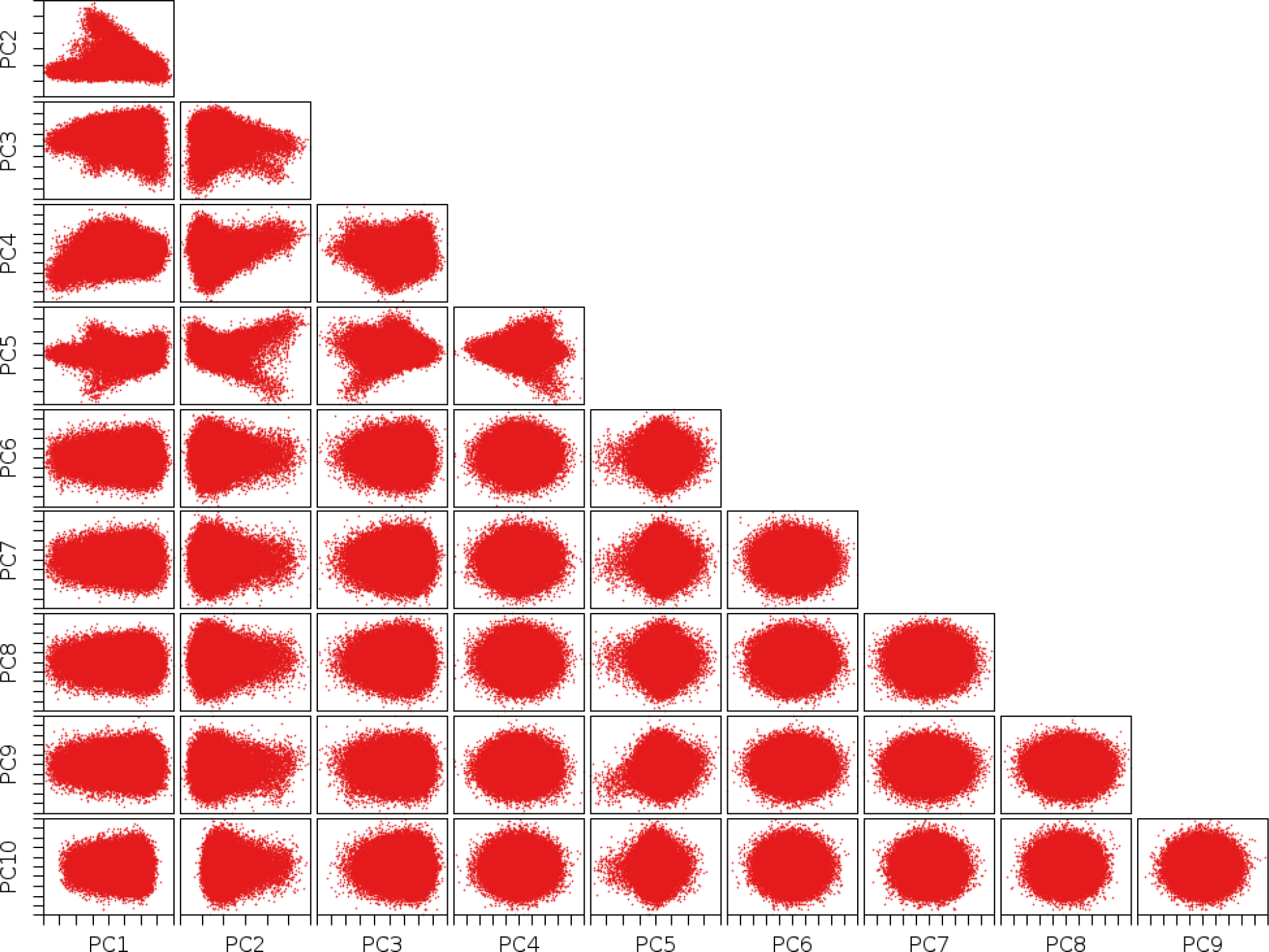

**Supplementary Figure 2 Results of PCA with k-means clustering for all PCs**

This is an expanded set of plots similar to Figure 1, except that plots of all pairs of top PCs are displayed.

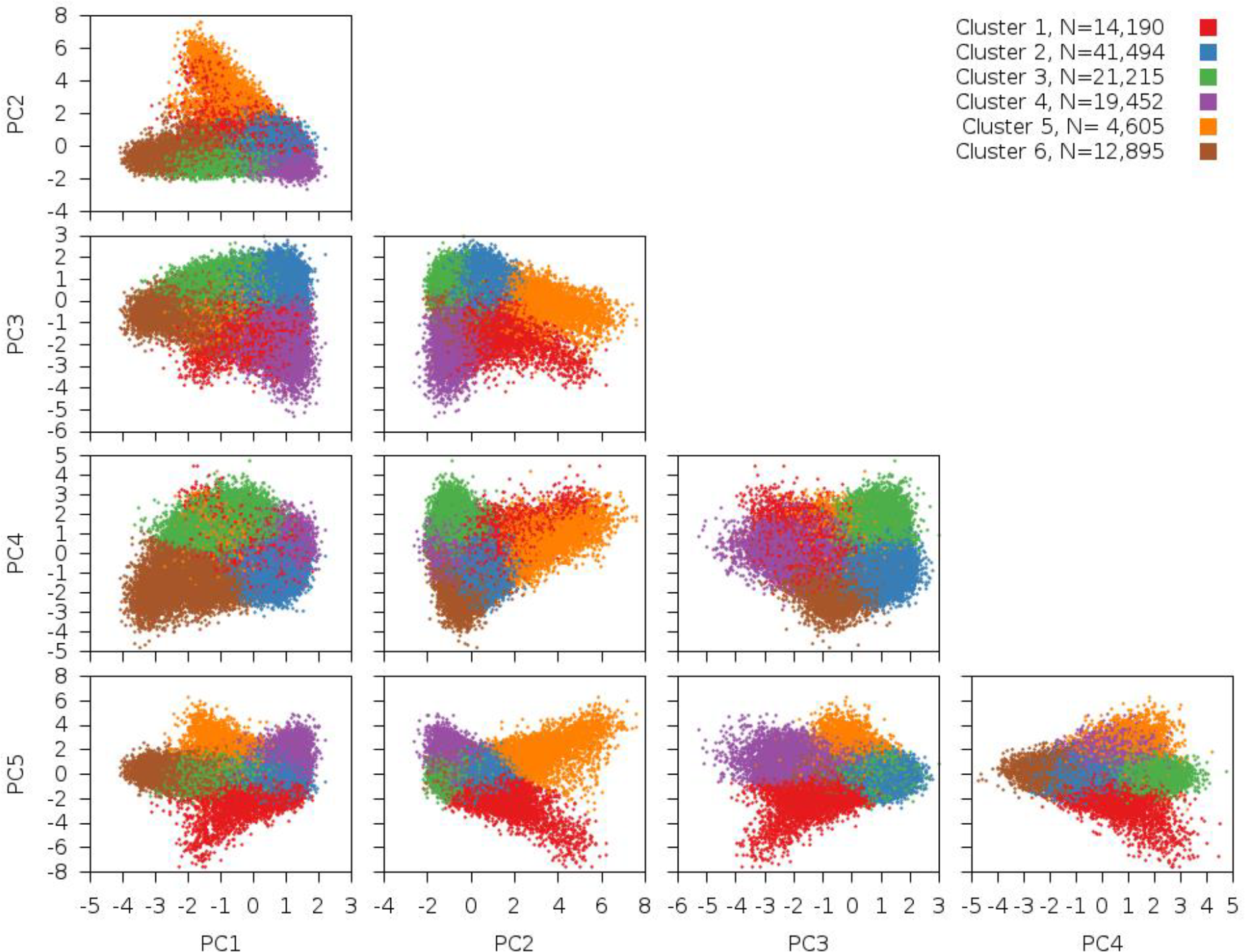

**Supplementary Figure 3 Results of PCA with projection of PoBI samples for all PCs**

This is an expanded set of plots similar to Figure 2, except that plots of all pairs of top PCs are displayed.

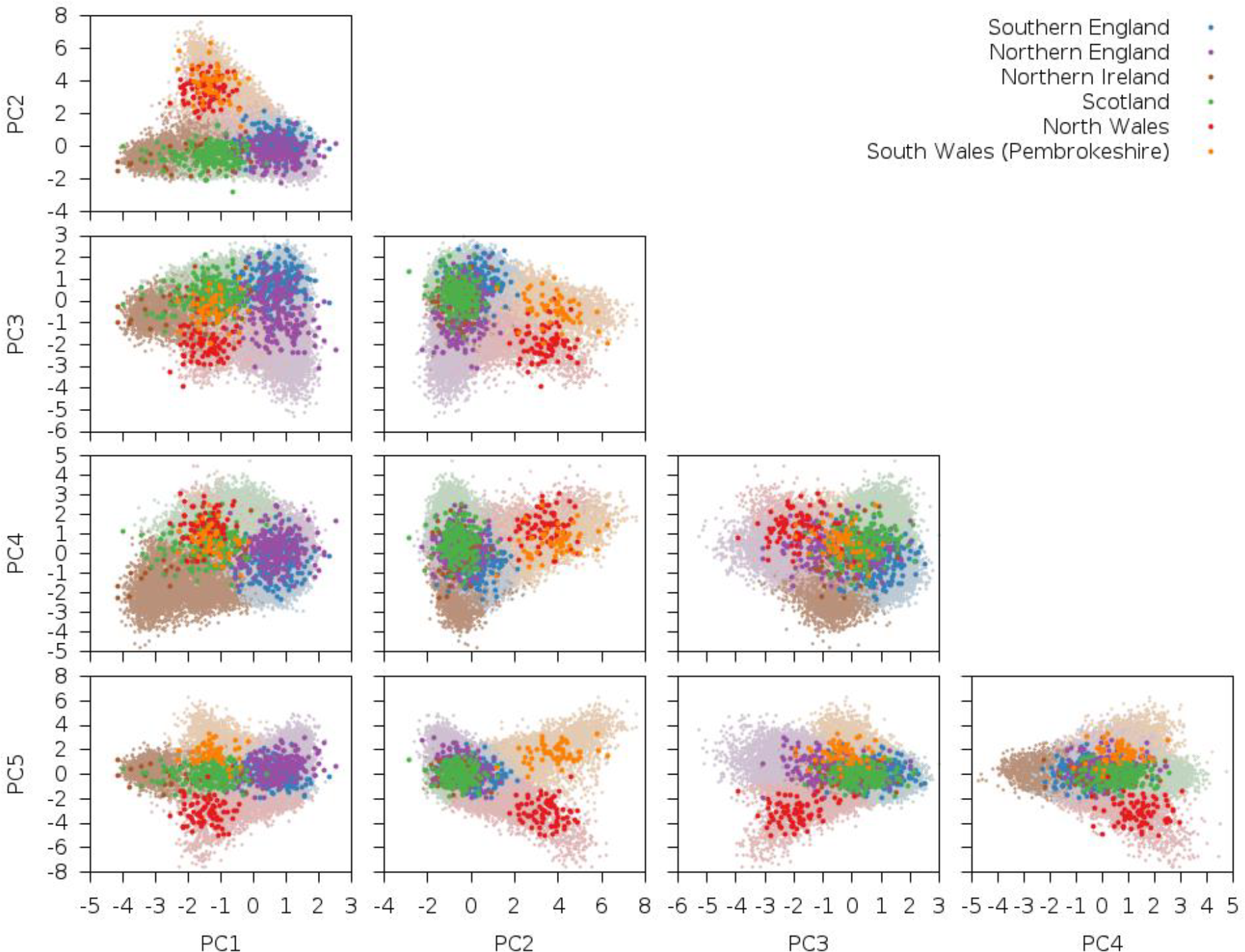

**Supplementary Figure 4 Results of PCA with projection of ancient samples for all PCs**

This is an expanded set of plots similar to Figure 3, except that plots of all pairs of top PCs are displayed.

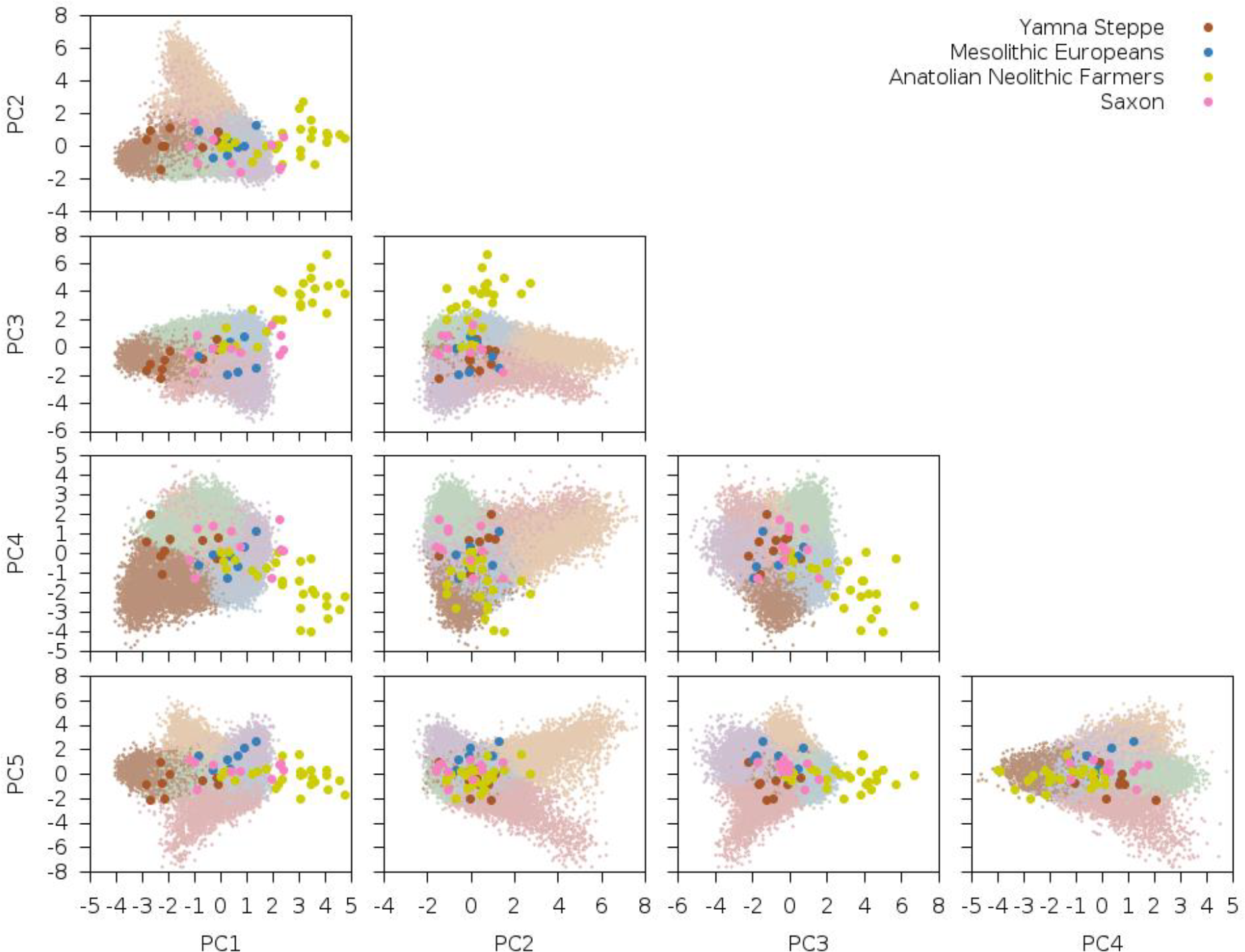

**Supplementary Figure 5 Results of PCA with projection of POPRES samples for all PCs**

This set of plots is similar to Supplementary Figure 2, except that POPRES samples are projected on top.

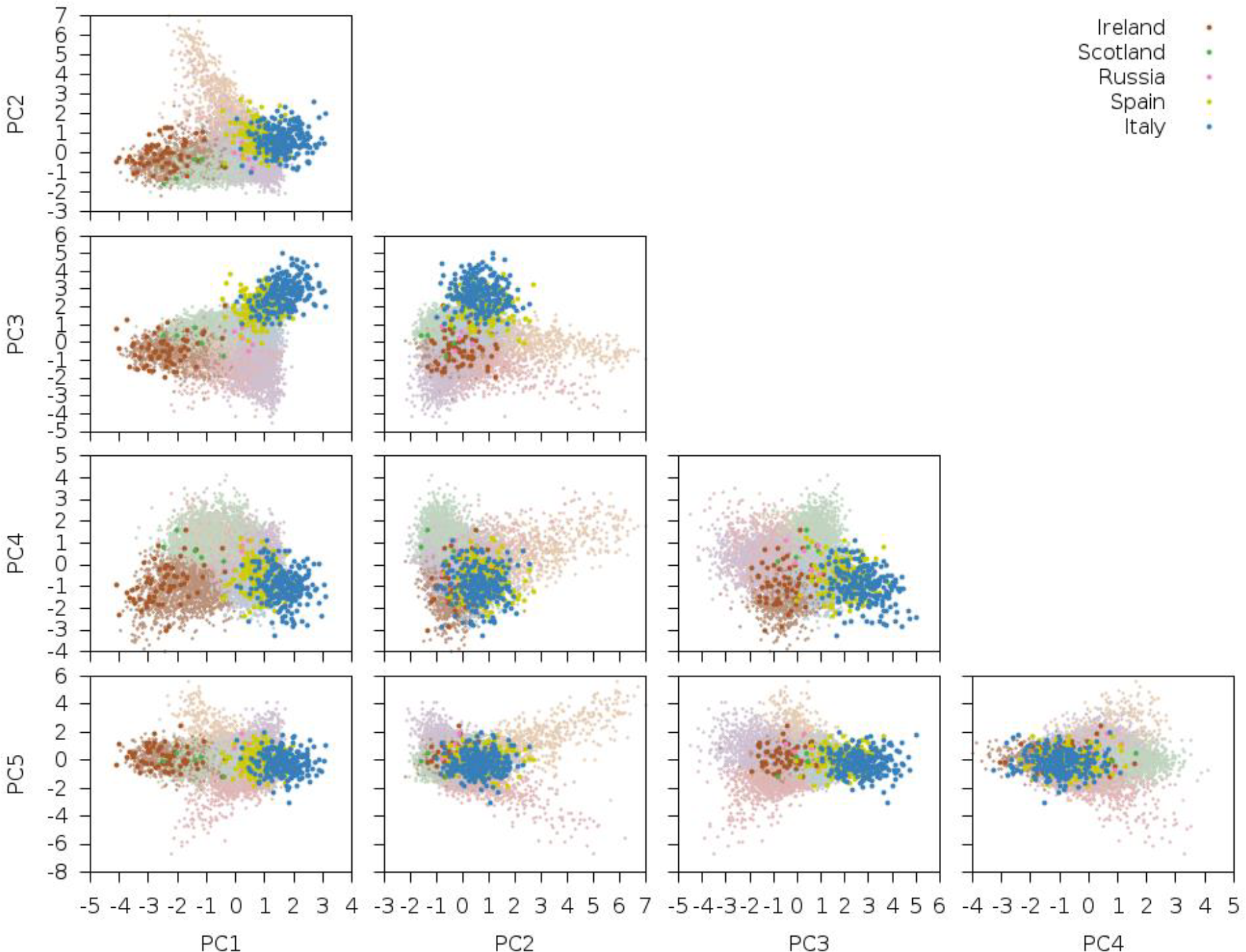

**Supplementary Figure 6 Selection statistic for UK Biobank along PC1-PC5**

This is an expanded set of plots similar to Figure 4, except that plots for each of the top 5 PCs are displayed.

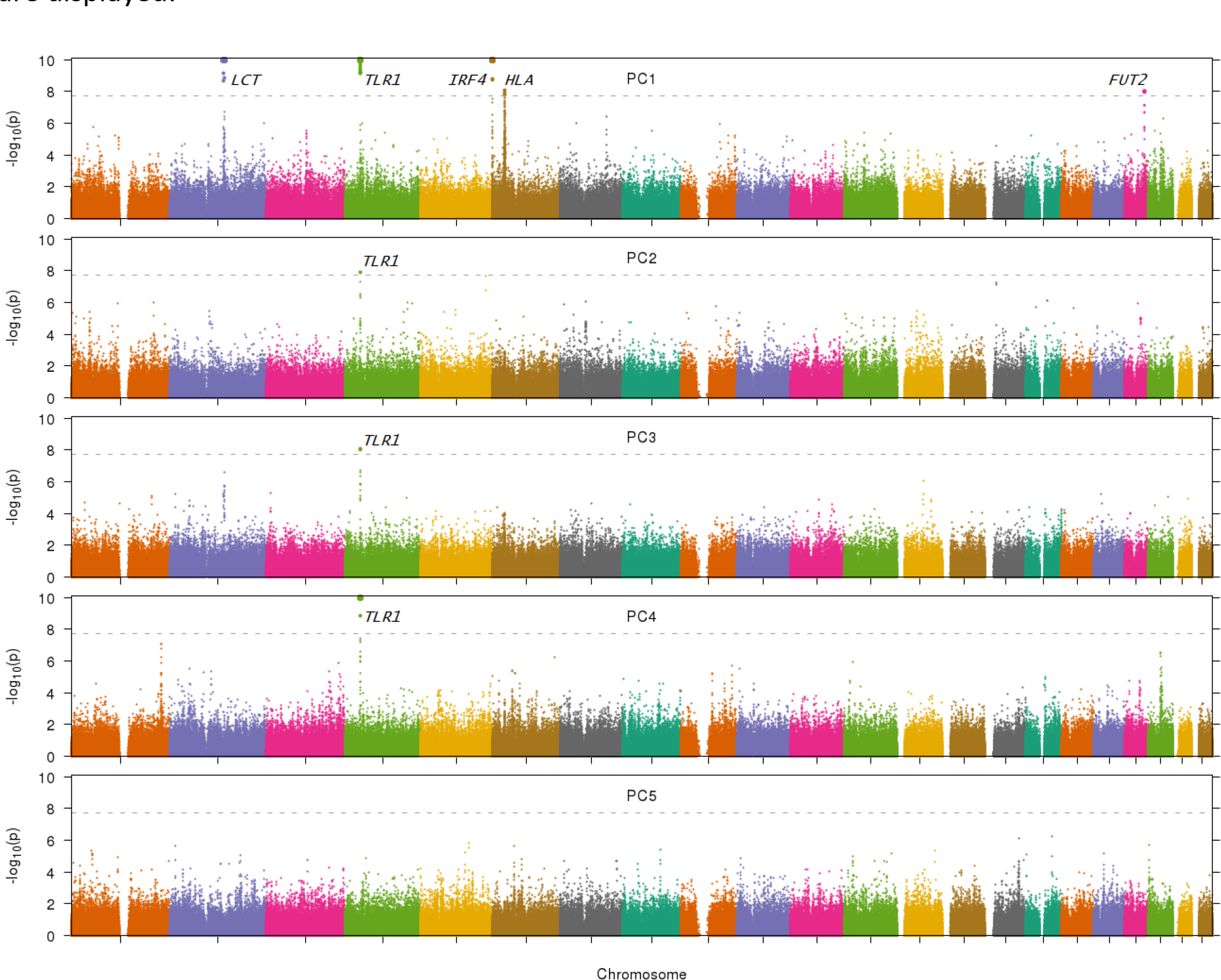

**Supplementary Figure 7 Results of initial PCA run**

This set of plots is similar to Supplementary Figure 1, except that long-range LD regions were not removed. Several of these PCs are dominated by regions of long-range LD. In particular, the three clusters along PC2 indicate 0, 1 or 2 copies of a chromosome 8 inversion variant.

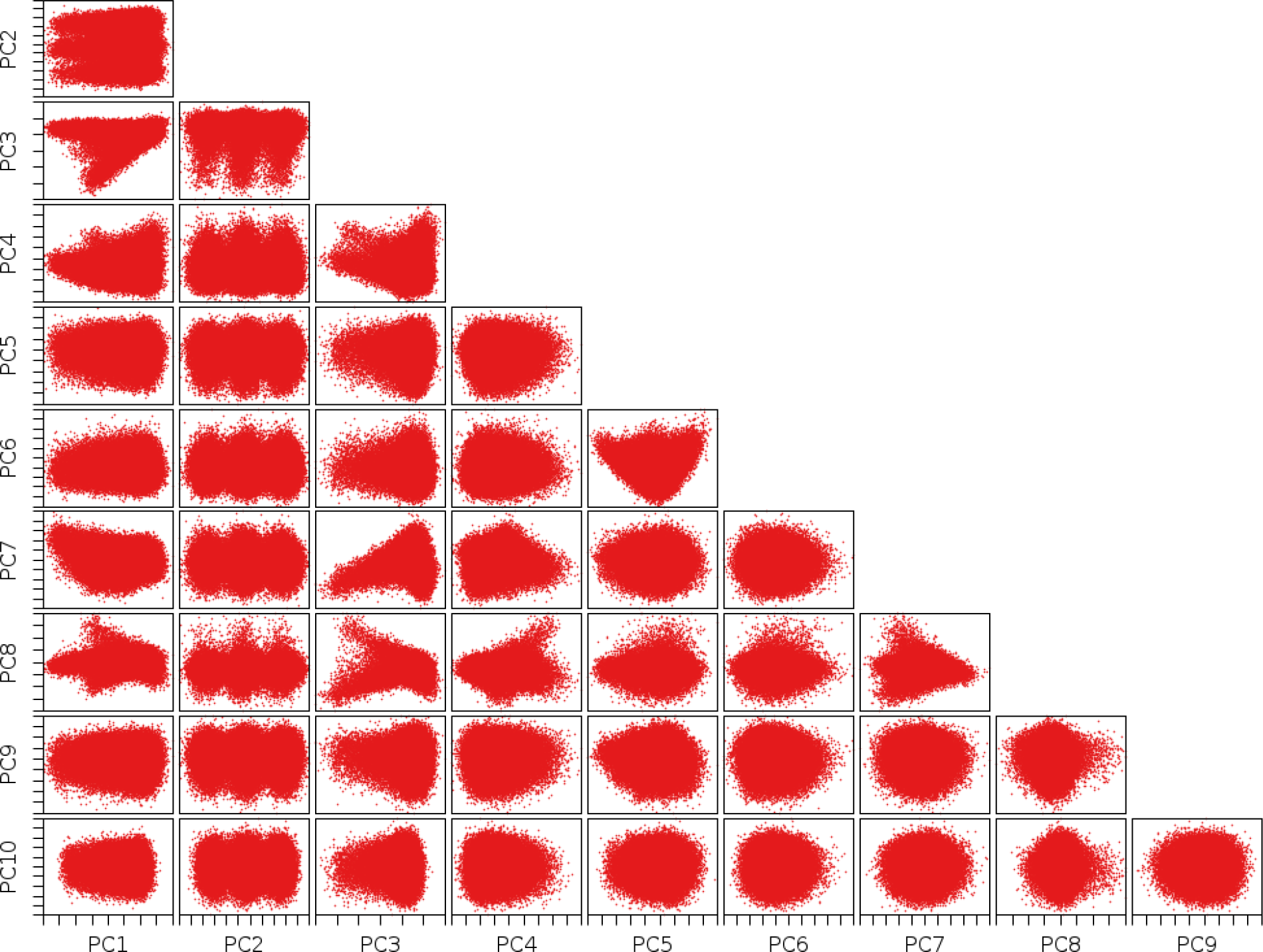

## Supplementary Tables

**Supplementary Table 1 PC eigenvalues and geographical correlations**

PC1-PC5 all had elevated eigenvalues, while PC6-PC10 had eigenvalues which were close to background levels. In the case where there are two equal-sized sample sets from distinct populations, the *F_ST_* between the two populations can be estimated from the top eigenvalue (*λ*) via the following formula: *F_ST_* = (*λ* − 1)/*N*, where *N* is the total number of samples. The top eigenvalue reflects an *F_ST_* of 1.76 × 10^−4^, indicating very subtle population structure within the UK. PC1 was most strongly correlated with east-west birth coordinate and PC2 was most strongly correlated with north-south birth coordinate.

**Table 1.**
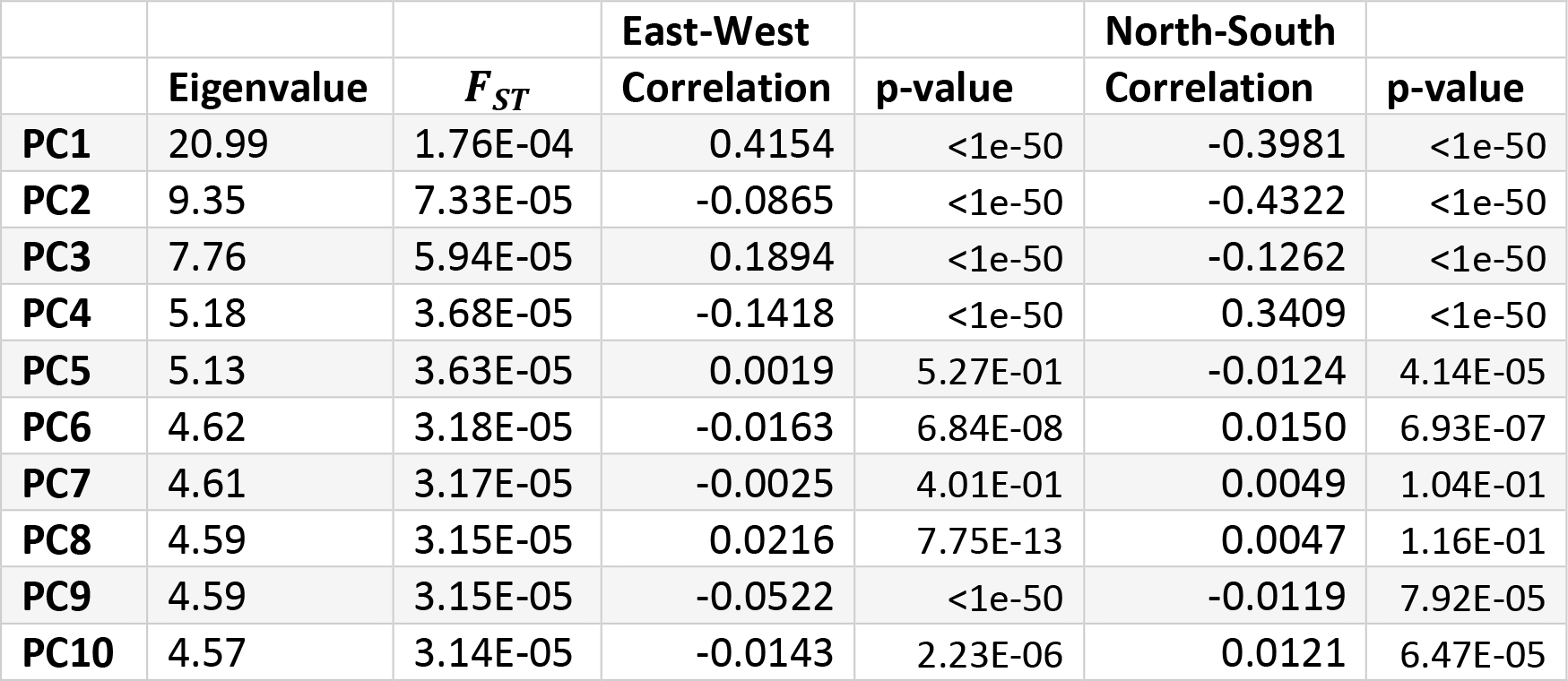

**Supplementary Table 2 Expanded results of *f*_4_ statistics in ancient and modern British samples**

We report *f*_4_ statistics of of the form *f*_4_(*Steppe*, *Neolithic Farmer*; *Pop*l, *Pop*2), representing a z-score with positive values indicating more Steppe ancestry in *Pop1* than *Pop2*. Samples for *Pop1* were either modern Celtic (Scotland and Wales) or ancient Saxon. Samples for *Pop2* were modern Anglo-Saxon (southern and eastern England).

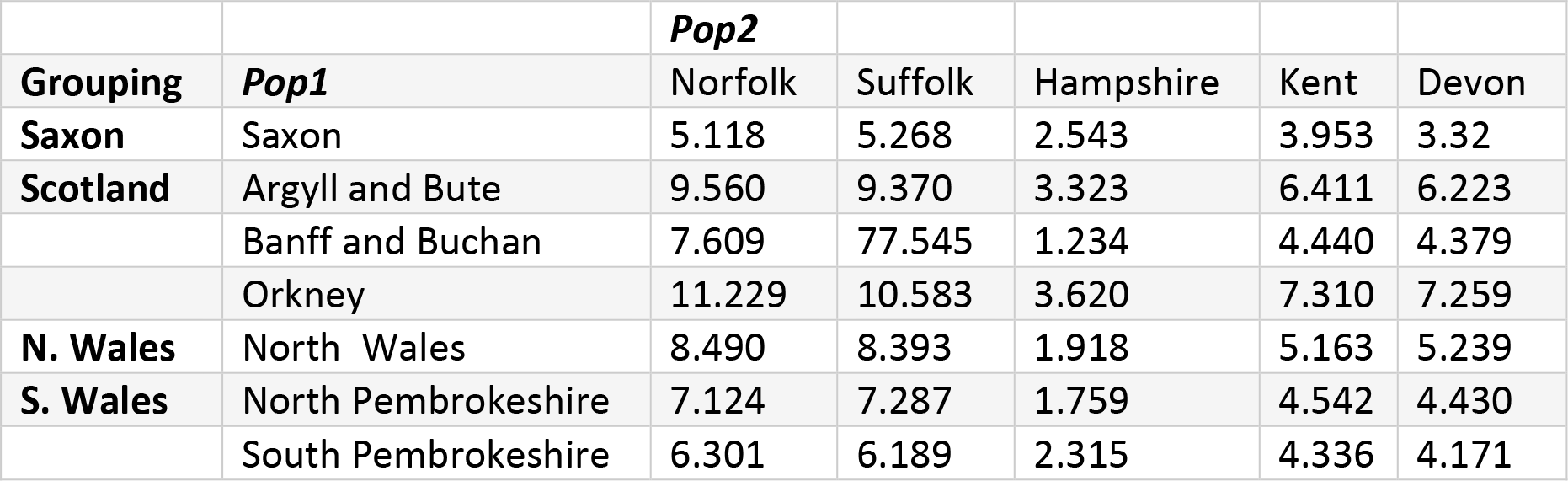

|  |  | ***Pop2*** |  |  |  |  |
| --- | --- | --- | --- | --- | --- | --- |
| **Grouping** | ***Pop1*** | Norfolk | Suffolk | Hampshire | Kent | Devon |
| **Saxon** | Saxon | 5.118 | 5.268 | 2.543 | 3.953 | 3.32 |
| **Scotland** | Argyll and Bute | 9.560 | 9.370 | 3.323 | 6.411 | 6.223 |
|  | Banff and Buchan | 7.609 | 77.545 | 1.234 | 4.440 | 4.379 |
|  | Orkney | 11.229 | 10.583 | 3.620 | 7.310 | 7.259 |
| **N. Wales** | North Wales | 8.490 | 8.393 | 1.918 | 5.163 | 5.239 |
| **S. Wales** | North Pembrokeshire | 7.124 | 7.287 | 1.759 | 4.542 | 4.430 |
|  | South Pembrokeshire | 6.301 | 6.189 | 2.315 | 4.336 | 4.171 |

**Supplementary Table 3 Suggestive signals of selection in UK Biobank**

We report the top signal of natural selection for each locus not reaching genome-wide significance (*p* > 1.96 × 10^−8^) but yielding a suggestive signal (*p* < 1.00 × 10^−6^) along any of the top five PCs. Neighboring SNPs <1Mb apart with suggestives significant signals were grouped together into a single locus.

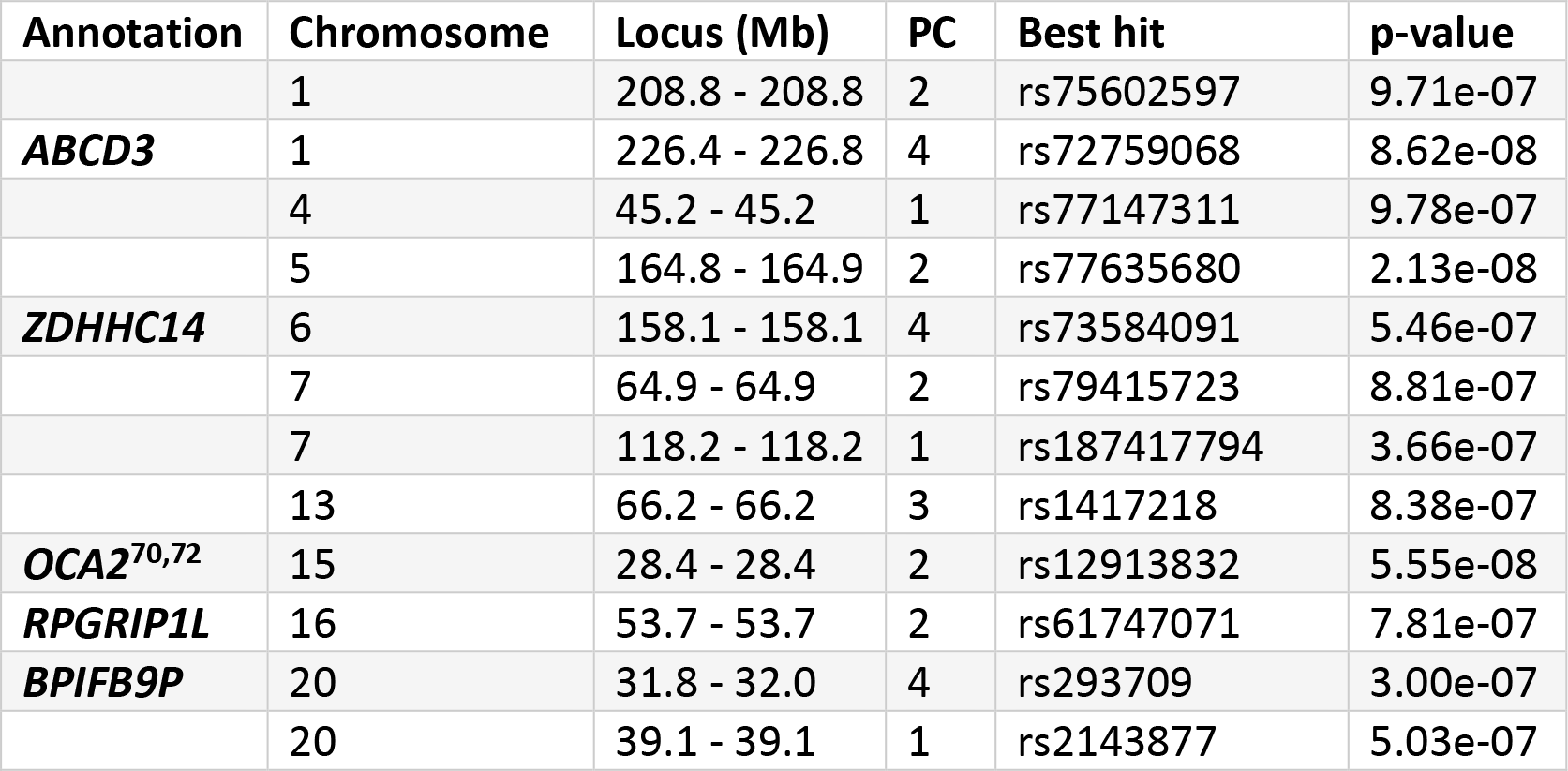

**Supplementary Table 4 Allele frequency of FUT2 alleles**

We report the allele frequency of the most significant hit, rs601338, along with two other linked SNPs in GERA and the PoBI datasets.

| **Dataset** | **Cluster** | **rs601338**<br>**(G/A)** | **rs492602**<br>**(G/A)** | **rs676388**<br>**(T/C)** |
| --- | --- | --- | --- | --- |
| **UK Biobank** | Northern Ireland | 0.4406 | 0.4413 | 0.4207 |
|  | Northeast England | 0.4633 | 0.4638 | 0.4407 |
|  | Pembrokeshire | 0.4864 | 0.4871 | 0.4580 |
|  | North Wales | 0.5006 | 0.501 | 0.4781 |
|  | Yorkshire | 0.5025 | 0.503 | 0.4763 |
|  | East Anglia | 0.5109 | 0.5111 | 0.4847 |
| **GERA** | Irish |  | 0.4754 | 0.4522 |
|  | Northern European |  | 0.5215 | 0.4962 |
|  | Southern European |  | 0.5248 | 0.5021 |
|  | Ashkenazi Jewish |  | 0.5530 | 0.5200 |
|  | Eastern European |  | 0.5840 | 0.5586 |
| **PoBI** | Argyll and Bute |  |  | 0.2949 |
|  | North Pembrokeshire |  |  | 0.3171 |
|  | Banff and Buchan |  |  | 0.3365 |
|  | Northern Ireland |  |  | 0.3667 |
|  | Cumbria |  |  | 0.4039 |
|  | Derbyshire |  |  | 0.4091 |
|  | Dorset |  |  | 0.4125 |
|  | Herefordshire |  |  | 0.4259 |
|  | Worcestershire |  |  | 0.4265 |
|  | Lancashire |  |  | 0.4306 |
|  | Devon |  |  | 0.4416 |
|  | Yorkshire |  |  | 0.4517 |
|  | Orkney |  |  | 0.4531 |
|  | Lincolnshire |  |  | 0.4619 |
|  | Kent |  |  | 0.4661 |
|  | Suffolk |  |  | 0.4699 |
|  | Cornwall |  |  | 0.4716 |
|  | Leicestershire |  |  | 0.4726 |
|  | North Wales |  |  | 0.4737 |
|  | Northeast England |  |  | 0.4844 |
|  | Cheshire |  |  | 0.4875 |
|  | Forest of Dean |  |  | 0.4881 |
|  | Norfolk |  |  | 0.4951 |
|  | Nottinghamshire |  |  | 0.5000 |
|  | Northamptonshire |  |  | 0.5000 |
|  | Oxfordshire |  |  | 0.5054 |
|  | Sussex |  |  | 0.5167 |
|  | Gloucestershire |  |  | 0.5417 |
|  | South Pembrokeshire |  |  | 0.5417 |
|  | Hampshire |  |  | 0.5833 |

**Supplementary Table 5 Discrete test for natural selection at *FUT2* in GERA**

We report results of tests for selection using discrete subpopulations for *FUT2* in the GERA data set. *FUT2* does not reach genome-wide significance in the GERA dataset, however there are several suggestive signals when comparing the “Irish” subgroup with the Northern European and Eastern European subpopulation.

| $F_{ST}$ | EE | IR | NE | SE |
| --- | --- | --- | --- | --- |
| AJ | 6.84E-03 | 6.71E-03 | 6.54E-03 | 3.45E-03 |
| EE |  | 9.44E-04 | 7.23E-04 | 2.39E-03 |
| IR |  |  | 1.26E-04 | 1.91E-03 |
| NE |  |  |  | 1.80E-03 |

| rs492602 | EE | IR | NE | SE |
| --- | --- | --- | --- | --- |
| AJ | 5.96E-01 | 1.84E-01 | 5.83E-01 | 5.05E-01 |
| EE |  | 1.48E-06 | 1.58E-03 | 9.25E-02 |
| IR |  |  | 1.34E-07 | 1.16E-01 |
| NE |  |  |  | 9.12E-01 |

| rs676388 | EE | IR | NE | SE |
| --- | --- | --- | --- | --- |
| AJ | 5.13E-01 | 2.45E-01 | 6.79E-01 | 6.73E-01 |
| EE |  | 2.49E-06 | 1.69E-03 | 1.10E-01 |
| IR |  |  | 4.53E-07 | 1.12E-01 |
| NE |  |  |  | 8.46E-01 |

**Supplementary Table 6 Discrete test for natural selection at *FUT2* in UK**

Biobank We report results of tests for selection using discrete subpopulations for *FUT2* in the UK Biobank data set, using the UK Biobank subpopulations derived from *k*-means clustering. With 15 comparisons per SNP and 510,665 SNPs (p-value threshold of 6.53 × 10^−9^), we are still able to find genome-wide-significant results when comparing East Anglia with Northeast England as well as Northern Ireland.

| $F_{ST}$ | 2 | 3 | 4 | 5 | 6 |
| --- | --- | --- | --- | --- | --- |
| 1 | 7.46E-05 | 1.23E-04 | 8.53E-05 | 2.60E-04 | 2.23E-04 |
| 2 |  | 1.10E-04 | 7.36E-05 | 2.90E-04 | 2.67E-04 |
| 3 |  |  | 1.48E-04 | 3.10E-04 | 1.22E-04 |
| 4 |  |  |  | 3.75E-04 | 2.96E-04 |
| 5 |  |  |  |  | 3.43E-04 |

| rs601338 | 2 | 3 | 4 | 5 | 6 |
| --- | --- | --- | --- | --- | --- |
| 1 | 1.42E-01 | 1.95E-05 | 7.94E-01 | 2.72E-01 | 1.42E-07 |
| 2 |  | 2.61E-09 | 2.22E-01 | 6.45E-02 | 6.04E-09 |
| 3 |  |  | 2.38E-05 | 9.12E-02 | 9.31E-03 |
| 4 |  |  |  | 2.77E-01 | 1.28E-06 |
| 5 |  |  |  |  | 1.47E-03 |

| rs492602 | 2 | 3 | 4 | 5 | 6 |
| --- | --- | --- | --- | --- | --- |
| 1 | 1.51E-01 | 1.65E-05 | 8.49E-01 | 2.72E-01 | 1.65E-07 |
| 2 |  | 3.11E-09 | 2.29E-01 | 6.86E-02 | 7.35E-09 |
| 3 |  |  | 2.50E-05 | 8.93E-02 | 9.86E-03 |
| 4 |  |  |  | 2.85E-01 | 1.42E-06 |
| 5 |  |  |  |  | 1.48E-03 |

| rs676388 | 2 | 3 | 4 | 5 | 6 |
| --- | --- | --- | --- | --- | --- |
| 1 | 2.91E-01 | 1.30E-05 | 9.82E-01 | 1.97E-01 | 3.28E-07 |
| 2 |  | 3.38E-08 | 2.13E-01 | 4.27E-02 | 1.07E-07 |
| 3 |  |  | 1.25E-04 | 2.06E-01 | 2.12E-02 |
| 4 |  |  |  | 2.17E-01 | 1.31E-05 |
| 5 |  |  |  |  | 9.38E-03 |

**Supplementary Table 7 Independence of UK Biobank and ancient Eurasian scans for selection**

The UK Biobank and ancient Eurasian selection statistics were not inflated genome-wide, nor at the overlapping SNPs in both datasets. Similarly, the combined selection statistic was not inflated either. The correlation between the two statistics is also small, with the UK Biobank PC1 and ancient Eurasian statistics being most correlated with *r* = 0.188.

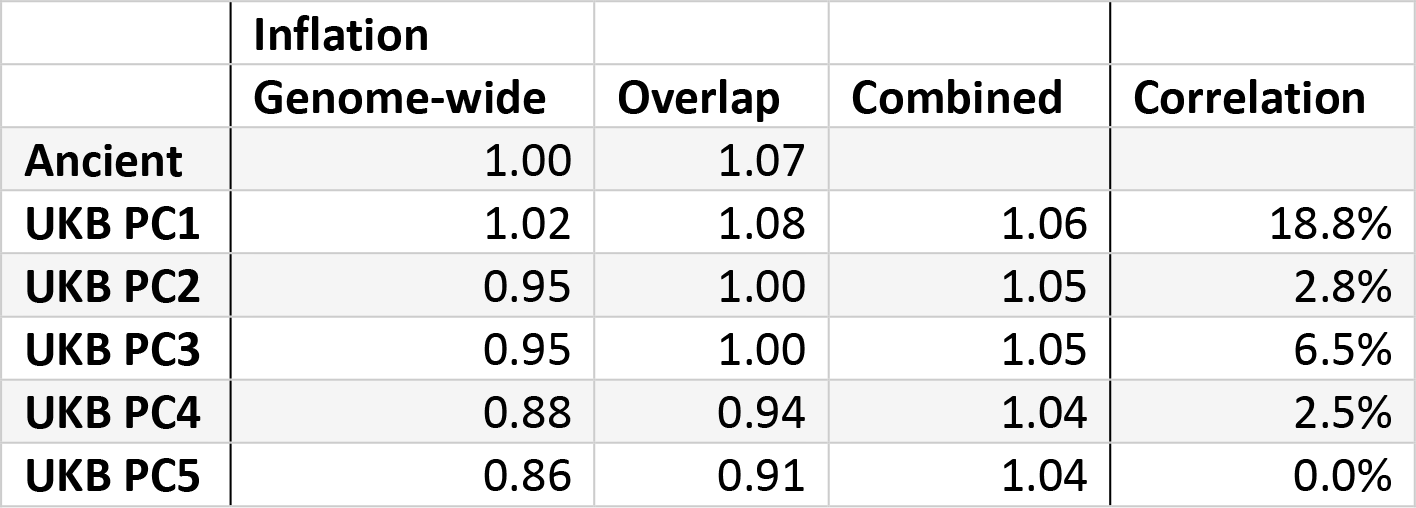

**Supplementary Table 8 Phenotype associations at SNPs with signals of selection**

We tested SNPs with genome-wide significant signals of selection in the constituent UK Biobank or ancient Eurasian scans or the combined scan for association with 15 phenotypes in the UK Biobank data set, using the top 5 PCs as covariates.

| Locus | Chr | Pos (Mb) | Phenotype | Top SNP | p-value |
| --- | --- | --- | --- | --- | --- |
| **LCT** | 2 | 134.9 - 137.0 | lung_FVCzSMOKE | rs6716536 | 4.90e-10 |
| **TLR1** | 4 | 38.8 - 38.9 | disease_ALLERGY_ECZEMA_DIAGNOSED | rs5743614 | 5.80e-26 |
| **SLC22A4** | 5 | 131.4 - 131.8 | body_HEIGHTz | rs1050152 | 3.70e-18 |
|  |  |  | disease_ASTHMA_DIAGNOSED | rs2188962 | 6.00e-09 |
| **F12** | 5 | 176.8 - 176.8 | body_HEIGHTz | rs2545801 | 4.80e-11 |
| **HLA** | 6 | 28.3 - 33.0 | body_HEIGHTz | rs2256183 | 4.10e-23 |
|  |  |  | body_WHRadjBMIz | rs521977 | 1.80e-10 |
|  |  |  | bp_DIASTOLICadjMEDz | rs521977 | 2.00e-10 |
|  |  |  | disease_ALLERGY_ECZEMA_DIAGNOSED | rs3135377 | 1.60e-11 |
|  |  |  | disease_ASTHMA_DIAGNOSED | rs204993 | 5.10e-09 |
|  |  |  | lung_FEV1FVCzSMOKE | rs3891175 | 3.50e-21 |
|  |  |  | lung_FVCzSMOKE | rs6456834 | 3.70e-09 |
| **FADS1** | 11 | 61.5 - 61.6 | bmd_HEEL_TSCOREz | rs174548 | 1.20e-08 |
| **ATXN2/SH2B3** | 12 | 111.9 - 112.9 | bp_DIASTOLICadjMEDz | rs3184504 | 8.00e-33 |
|  |  |  | bp_SYSTOLICadjMEDz | rs3184504 | 1.90e-13 |
|  |  |  | disease_HYPERTENSION_DIAGNOSED | rs3184504 | 1.30e-09 |
| **CYP1A2/CSK** | 15 | 75.0 - 75.1 | bp_DIASTOLICadjMEDz | rs2472304 | 1.10e-19 |
|  |  |  | bp_SYSTOLICadjMEDz | rs2472304 | 4.20e-10 |
|  |  |  | disease_HYPERTENSION_DIAGNOSED | rs2472304 | 2.60e-09 |

**Supplementary Table 9 PC-phenotype associations in UK Biobank**

We report the results of tests of associations (p-value) between top PCs and 15 phenotypes in UK Biobank. This analysis does not distinguish between environmental and genetic effects.

| Phenotype | PC1 | PC2 | PC3 | PC4 | PC5 |
| --- | --- | --- | --- | --- | --- |
| **bmd_HEEL_TSCOREz** | 2.33E-01 | 4.75E-01 | 2.39E-23 | 3.54E-07 | 2.97E-01 |
| **body_BMIz** | 2.36E-27 | 7.98E-01 | 2.59E-11 | 2.21E-03 | 1.14E-01 |
| **body_HEIGHTz** | <1e-50 | 4.66E-01 | <1e-50 | 8.38E-03 | 2.41E-03 |
| **body_WHRadjBMIz** | 2.75E-06 | 3.35E-01 | 3.71E-03 | 2.94E-24 | 1.06E-01 |
| **bp_DIASTOLICadjMEDz** | 7.34E-01 | 5.42E-01 | 2.70E-08 | 6.16E-13 | 2.37E-01 |
| **bp_SYSTOLICadjMEDz** | 1.75E-03 | 7.67E-02 | 6.31E-01 | 6.15E-07 | 3.09E-02 |
| **cov_EDU_COLLEGE** | 6.73E-01 | 9.75E-25 | 2.01E-29 | 8.59E-36 | 7.96E-07 |
| **cov_SMOKING_STATUS** | 7.13E-01 | 5.67E-01 | 3.61E-03 | 9.63E-07 | 8.08E-01 |
| **disease_ALLERGY_ECZEMA_DIAGNOSED** | 1.76E-16 | 1.50E-03 | 1.07E-08 | 7.15E-03 | 7.87E-01 |
| **disease_ASTHMA_DIAGNOSED** | 7.04E-01 | 2.50E-06 | 6.12E-01 | 2.89E-05 | 3.84E-01 |
| **disease_HYPERTENSION_DIAGNOSED** | 7.84E-01 | 2.45E-01 | 1.09E-03 | 5.92E-01 | 3.32E-01 |
| **lung_FEV1FVCzSMOKE** | 9.85E-01 | 3.31E-07 | 1.89E-13 | 5.58E-10 | 3.43E-07 |
| **lung_FVCzSMOKE** | 8.69E-02 | 1.93E-45 | 2.21E-03 | 3.48E-40 | 3.99E-04 |
| **repro_MENARCHE_AGE** | 1.14E-04 | 1.11E-05 | 1.93E-03 | 3.11E-01 | 3.75E-02 |
| **repro_MENOPAUSE_AGE** | 1.46E-15 | 7.51E-04 | 9.54E-07 | 1.27E-03 | 1.93E-01 |

**Supplementary Table 10 Significant or suggestive signals of selection in initial PCA run**

We report significant or suggestive signals of selection in the initial PCA run. Neighboring SNPs <1Mb apart with genome-wide significant signals were grouped together into a single locus. The significant signals may represent either signals of selection or regions of long-range LD. All of these regions were removed from the main PCA run (see Online Methods).

| Chrom | Locus (Mb) | PC | Best hit | *p*-value |
| --- | --- | --- | --- | --- |
| 1 | 2.2 - 2.2 | 3 | rs79907870 | 4.68E-07 |
| 1 | 54.8 - 54.8 | 1 | rs17390412 | 9.13E-07 |
| 1 | 56.0 - 56.1 | 8 | rs1875068 | 1.86E-08 |
| 2 | 88.7 - 88.7 | 4 | rs1713939 | 1.02E-07 |
| 2 | 133.3 - 144.0 | 1 | rs7570971 | 7.21E-18 |
|  |  | 4 | rs1446585 | 3.02E-14 |
|  |  | 7 | rs1446585 | 3.09E-13 |
|  |  | 10 | rs72847650 | <1e-50 |
| 2 | 159.8 - 160.0 | 7 | rs1522699 | 3.89E-07 |
| 2 | 223.9 - 223.9 | 7 | rs1900725 | 2.71E-07 |
| 3 | 46.3 - 46.4 | 7 | rs9990343 | 2.80E-08 |
| 4 | 23.3 - 23.3 | 3 | rs114557362 | 2.95E-07 |
| 4 | 38.7 - 38.9 | 1 | rs4833095 | 9.29E-16 |
|  |  | 3 | rs4833095 | 8.54E-11 |
|  |  | 4 | rs4833095 | 1.21E-10 |
|  |  | 7 | rs4833095 | 2.18E-16 |
| 5 | 60.6 - 60.6 | 3 | rs10471511 | 6.04E-07 |
| 5 | 101.5 - 101.6 | 8 | rs411954 | 1.20E-08 |
| 5 | 114.8 - 114.8 | 8 | rs895291 | 5.87E-07 |
| 5 | 164.8 - 164.9 | 3 | rs77635680 | 6.70E-10 |
| 6 | 0.4 - 0.7 | 1 | rs62389423 | 1.29E-47 |
|  |  | 7 | rs62389423 | 1.57E-09 |
| 6 | 23.9 - 36.7 | 1 | rs151341075 | 3.76E-11 |
|  |  | 3 | rs2253908 | 5.43E-07 |
|  |  | 4 | rs151341075 | 3.35E-17 |
|  |  | 5 | rs3131618 | <1e-50 |
|  |  | 6 | rs204999 | <1e-50 |
|  |  | 7 | rs2596573 | 3.27E-54 |
|  |  | 8 | rs41268932 | 2.58E-23 |
|  |  | 9 | rs2596573 | <1e-50 |
|  |  | 10 | rs9266258 | 4.14E-07 |
| 6 | 46.8 - 46.8 | 7 | rs9395218 | 9.92E-07 |
| 6 | 86.0 - 87.0 | 10 | rs2816583 | 2.23E-08 |
| 7 | 41.4 - 41.4 | 1 | rs76920365 | 3.66E-07 |
| 7 | 64.4 - 66.4 | 3 | rs79415723 | 1.17E-08 |
| 7 | 118.2 - 118.3 | 1 | rs187417794 | 1.13E-07 |
| 8 | 7.2 - 12.7 | 2 | rs11250099 | <1e-50 |
| 9 | 14.0 - 14.0 | 3 | rs12380860 | 4.45E-07 |
| 10 | 133.2 - 133.2 | 4 | rs57105422 | 7.45E-07 |
| 15 | 28.4 - 28.4 | 3 | rs12913832 | 9.17E-10 |
| 15 | 50.8 - 50.8 | 5 | rs148783236 | 3.18E-194 |
|  |  | 6 | rs148783236 | <1e-50 |
|  |  | 8 | rs148783236 | 2.18E-13 |
|  |  | 9 | rs148783236 | <1e-50 |
| 16 | 9.5 - 9.5 | 4 | rs12149526 | 6.26E-07 |
| 16 | 26.5 - 26.5 | 3 | rs73528772 | 4.01E-07 |
| 16 | 53.7 - 53.7 | 3 | rs61747071 | 1.36E-07 |
| 16 | 89.7 - 89.8 | 4 | rs449882 | 5.37E-07 |
| 17 | 29.6 - 29.6 | 3 | rs11655238 | 5.98E-07 |
| 19 | 33.8 - 33.8 | 3 | rs41355649 | 3.65E-08 |
| 19 | 49.2 - 50.2 | 1 | rs601338 | 1.05E-09 |
| 20 | 39.1 - 39.1 | 1 | rs2143877 | 8.34E-08 |
| 22 | 32.9 - 32.9 | 5 | rs115815765 | 7.40E-31 |
|  |  | 6 | rs115815765 | <1e-50 |

